# SIMORGH: Ensemble-Aware Geometric Deep Learning for Apo-State and Cryptic Ligand Binding Site Prediction

**DOI:** 10.64898/2026.10.02.756215

**Authors:** Omid Mokhtari, Amirhossein Naghsh Nilchi, Björn Grüning, Yasaman Karami, Hamed Khakzad

**Affiliations:** Université de Lorraine, Inria, F-54000 Nancy, France; Chair for Bioinformatics, Department of Computer Science, University of Freiburg, Georges-Koehler-Allee 079, 79110 Freiburg, Germany; Institute of Experimental and Clinical Pharmacology and Toxicology, Faculty of Medicine, University of Freiburg, Freiburg, Germany

## Abstract

Most computational methods identify protein–ligand binding sites from ligand-bound (holo) protein structures, where the binding pocket is already preorganized. Although convenient for benchmarking, this setting differs from the practical drug discovery scenario, in which binding sites must be inferred from ligand-free (apo) proteins. In fact, binding-competent conformations may represent only a subset of the accessible structural ensemble, while transient and cryptic pockets can emerge through conformational fluctuations. Consequently, prediction from a single static structure is often insufficient. Here we present SIMORGH, an ensemble-aware SE(3)-equivariant geometric learning framework that predicts ligand binding sites by integrating information across protein conformational ensembles. SIMORGH employs a two-stage architecture consisting of a structure-level equivariant encoder that learns geometric representations from individual conformations and a lightweight ensemble-level aggregation module that combines them into residue-level binding predictions while scaling near-linearly with both protein and ensemble size. We evaluate SIMORGH across multiple complementary settings, including the PLINDER apo benchmark, molecular dynamics trajectories spanning more than 200 cryptic-pocket proteins, and established community benchmarks. SIMORGH consistently outperforms existing static and dynamics-aware methods under the challenging ligand-free setting, while identifying cryptic binding pockets, maintaining strong prediction consistency between paired apo and holo structures, and scaling efficiently to large multi-chain complexes. To facilitate reproducible, transparent, and accessible use, Simorgh is available through the Galaxy Europe server (https://usegalaxy.eu).

## 1 Introduction

Accurate prediction of protein–ligand binding sites is a central problem in structure-based drug discovery, directly influencing hit identification, lead optimization, and the rational design of selective therapeutics [1]. Despite remarkable advances in structural biology and geometric deep learning, most computational approaches continue to infer binding sites from a static protein structure. This assumption fundamentally conflicts with the dynamic nature of proteins, whose ligand-binding competent states can emerge only transiently within a broader conformational ensemble. Consequently, binding sites may remain partially formed or entirely absent in experimentally resolved apo structures, particularly for cryptic pockets and highly flexible binding interfaces [2]. These limitations are further amplified by current benchmarking practices, which predominantly evaluate methods on ligand-bound (holo) structures, where pocket geometry is already stabilized around the ligand and therefore provides substantially more structural information than is available in realistic prospective drug discovery [3]. As a result, performance on conventional benchmarks may not accurately reflect a model’s ability to identify ligand binding sites directly from ligand-free proteins. Moreover, recent studies have highlighted that structural similarity between training and evaluation datasets, particularly for modern structure prediction and co-folding pipelines, can further inflate benchmark performance and obscure true model generalization under realistic ligand-free prediction scenarios [4].

The increasing availability of large-scale molecular dynamics (MD) datasets [5–8] has created new opportunities for dynamics-aware machine learning approaches [9–11] capable of capturing conformational heterogeneity. However, integrating conformational ensembles into geometric learning remains challenging. Atomistic MD simulations generate large ensembles of conformations per protein, making end-to-end processing computationally prohibitive for learning frameworks. Furthermore, the high computational cost of MD simulations limits the scalability of generating training data for data-intensive GNN-based models. These constraints have motivated alternative strategies based on coarse-grained physical simulations [12, 13] and deep generative conformational samplers [14, 15], which substantially increase structural coverage while trading off some degree of physical realism. Together, these observations highlight the need for learning frameworks capable of exploiting large conformational datasets while remaining robust to the structural variability encountered in prospective drug discovery.

On the algorithmic side, binding-site prediction has evolved from grid-based convolutional neural networks [16–18] toward graph neural networks (GNNs) [19, 20] and geometric transformers [21], with more recent methods integrating protein language models to leverage evolutionary information [3, 22, 23]. A particularly important advance has been the adoption of SE(3)-equivariant neural networks, which move beyond purely scalar representations by learning higher-order geometric features that naturally respect the rotational and translational symmetries of molecular structures [24–27]. Such representations are especially advantageous for modeling anisotropic interactions, including hydrogen bonding and *π*–*π* stacking, that are difficult to capture using coarse-grained representations without extensive feature engineering. Existing equivariant architectures generally balance expressivity and efficiency through different design choices: lightweight models such as EGNNs [27–29] avoid higher-order representations to reduce computational cost, whereas tensor-product-based architectures [30, 31] explicitly operate on higher-order irreducible representations (Irreps) at substantially greater computational expense. Despite their strong performance in small-molecule systems, their extension to macromolecular targets remains computationally prohibitive. Moreover, existing equivariant architectures are predominantly designed for single-structure inputs, leaving the central challenge of learning directly from protein conformational ensembles unresolved.

To address these challenges, we introduce **SIMORGH** (**S**et-**I**ntegrated **M**ulti-**O**rder **R**epresentation of **G**eometric **H**eterogeneity), an ensemble-aware SE(3)-equivariant framework for ligand binding-site prediction directly from protein conformational ensembles. SIMORGH decouples geometric representation learning from ensemble reasoning by independently modeling local geometry within individual conformations as residue-level equivariant representations, and subsequently integrating conformational variability through a tensor-order-aware attention mechanism. This formulation enables efficient learning from large collections of ligand-free protein conformations while maintaining near-linear computational scaling with both protein size and ensemble size. Across conventional holo benchmarks, SIMORGH achieves state-of-the-art or competitive performance while demonstrating substantially improved generalization under realistic apo-state evaluations, including a large cryptic pocket benchmark comprising 207 apo proteins. These results highlight the importance of ensemble-aware geometric learning for realistic structure-based drug discovery.

## 2 Results

### 2.1 SIMORGH decouples geometric representation learning from ensemble reasoning

SIMORGH is an ensemble-aware *SE*(3)-equivariant architecture that learns ligand binding sites directly from protein conformational ensembles through a two-stage framework, where local geometric representation learning and ensemble aggregation are performed separately (Fig. 1a, Fig. S1). In the first stage, the model independently learns *multi-order representations* (MORs) for each conformation by encoding local geometric environments through equivariant message passing. Although learning is performed at residue resolution, the encoder preserves atomistic geometric information by explicitly incorporating backbone-derived side-chain orientations and local reference frames (Table S1). Trained under binding-site supervision, these MORs become transferable geometric descriptors that capture both the chemical identity and spatial organization of individual conformations, providing the interface between the two learning stages.

**Figure 1:**
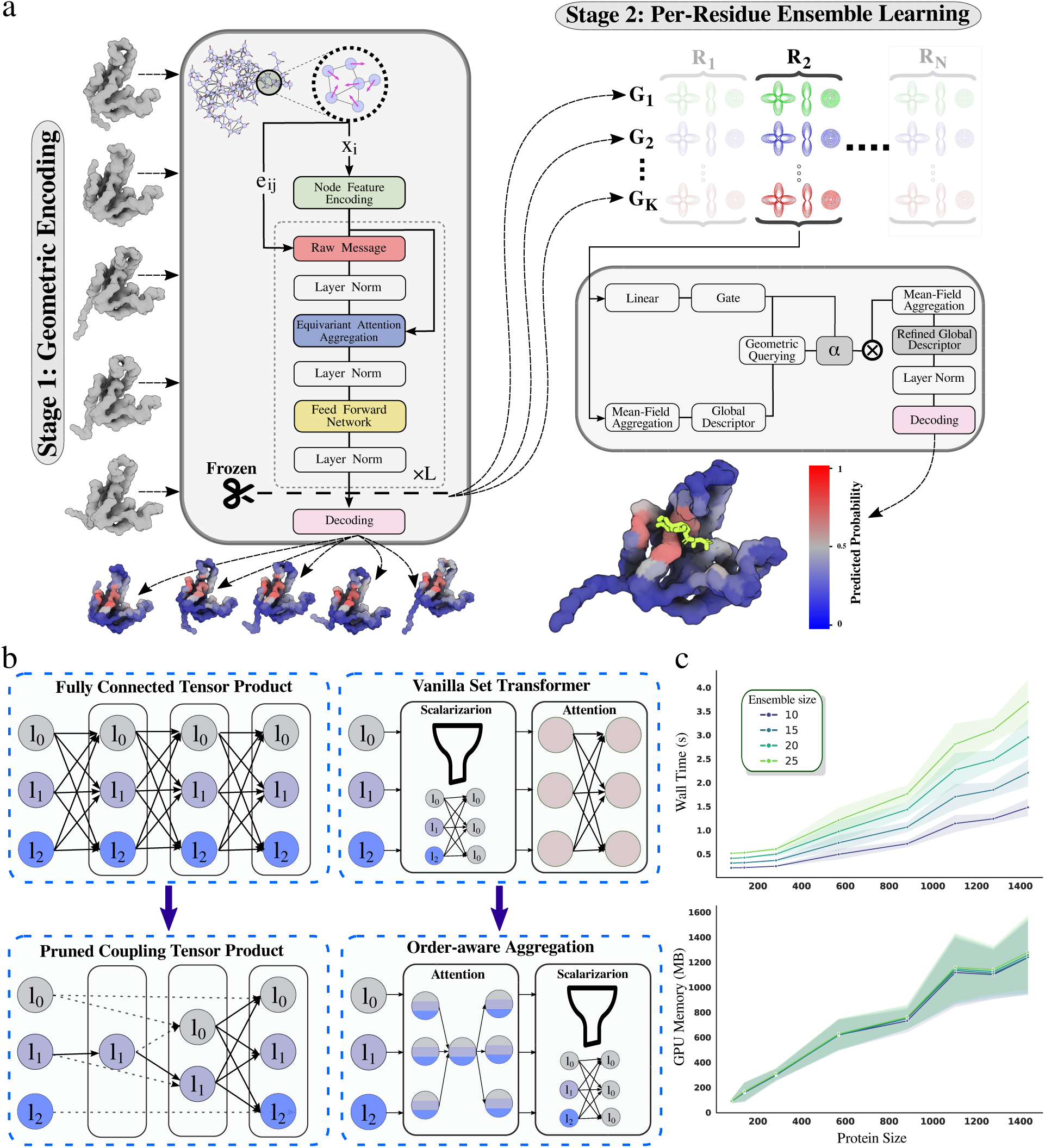
a: Overview of the SIMORGH network architecture. Stage 1 (Geometric Encoding) processes static structural inputs to extract latent MORs designed for transfer learning. Stage 2 (Per-Residue Ensemble Learning) aggregates these latent MORs across a conformational ensemble at the residue level to generate final predictions. b: Schematic representation of the architectural changes applied to standard message-passing layers. A fully connected tensor product (left) and vanilla set transformer aggregation (right) are modified into a pruned coupling tensor product and an order-aware aggregation module to yield a more lightweight, computationally efficient model. c: Inference computational requirement. Total wall time and peak GPU memory usage are evaluated as a function of protein size (ranging from 100 to 1,400 residues) across varying conformational ensemble sizes (10 to 25 conformations).

A central challenge in constructing expressive MORs is the reliance on higher-order irreps, whose computational cost increases sharply with tensor order. Rather than densely coupling all representation orders throughout message passing, SIMORGH introduces a channel-diversified communication strategy that factorizes message construction into successive interactions between neighboring residue representations, edge geometry, and the receiving node (Fig. 1b, Fig. S2). Vector channels (*ℓ* = 1) serve as the principal communication pathway, whereas second-order channels (*ℓ* = 2) are refined through lightweight residual interactions instead of repeated dense tensor products. Combined with the separate treatment of scalar and vector streams following previous equivariant architectures [21, 32], this substantially reduces computational complexity while preserving geometric expressiveness. Empirically, *ℓ*_2_ representations provide the best trade-off between predictive performance and computational efficiency (Supplementary Table S2).

In the second stage, MORs extracted from individual conformations are integrated through an order-aware ensemble aggregation module. Rather than applying conventional all-to-all attention across every pair of conformations, SIMORGH introduces a learned *Consensus Reference Token* that serves as a shared aggregation anchor (Fig. 1b). Each conformation communicates only with this consensus representation, reducing aggregation complexity from *O*(*K*^2^) to *O*(*K*) while maintaining independent attention across tensor orders. This formulation enables the model to selectively reinforce structural features that remain consistent across the ensemble while suppressing transient conformational fluctuations, allowing functional geometric signals to emerge from heterogeneous conformational landscapes.

Together, these design principles enable efficient equivariant learning over large conformational ensembles without sacrificing scalability. By combining channel-diversified message passing with consensus-based ensemble aggregation, SIMORGH exhibits near-linear scaling with both protein size and ensemble size while maintaining a peak GPU memory footprint that is largely independent of the number of conformations (Fig. 1c). In practice, evaluating a 1,400-residue protein complex across 25 conformations takes less than 4 seconds and requires approximately 1.3 GB of GPU memory, making ensemble-aware geometric learning practical for large-scale ligand-binding site prediction.

### 2.2 Learned ensemble aggregation consistently improves binding-site prediction across holo and apo settings

To quantify the contribution of ensemble modeling, we progressively evaluated four variants of our architecture, beginning from a static encoder and incrementally introducing ensemble representations and learned set aggregation (Table 1). We report results on both the similarity-filtered MISATO benchmark, derived from molecular dynamics trajectories, and the PLINDER apo benchmark, allowing us to assess whether the observed improvements generalize from holo-like to realistic ligand-free conditions.

**Table 1:** Progressive evaluation of the proposed architecture on holo and apo benchmarks. Performance is reported on the MISATO MD benchmark (holo) and the PLINDER benchmark (apo). The four variants progressively introduce the key components of SIMORGH: a static encoder, ensemble modeling, vanilla set transformer, and the proposed order-aware equivariant set aggregation. Bold indicates the best performance within each benchmark.

| Model Variant | Params<br>Overhead | MISATO (Holo) |  | PLINDER (Apo) |  |
| --- | --- | --- | --- | --- | --- |
|  |  | F1 | PR-AUC | F1 | PR-AUC |
| Static Encoder | 5.5M | 0.4194 | 0.4883 | 0.3735 | 0.4743 |
| + Ensemble Augmentation | – | 0.4484 | 0.5196 | 0.3819 | 0.4770 |
| + Learned Aggregation (SIMORGH) |  |  |  |  |  |
| Set Transformer | +186K | **0.4508** | **0.5314** | 0.3767 | **0.4973** |
| Order-Aware Consensus | +78K | 0.4455 | 0.5293 | **0.3920** | 0.4948 |

The *Static Encoder*, which processes a single protein conformation, consistently exhibited the weakest performance on both benchmarks, demonstrating the limitations of relying on a single structural snap-shot. Augmenting multiple conformations (*+ Ensemble*) substantially improved performance, confirming that conformational variability provides complementary information beyond any individual structure. This trend was remarkably consistent across both MISATO and PLINDER, despite their fundamentally different structural settings.

We next replaced simple ensemble integration with a learned aggregation module. To this end, we evaluated both the standard Set Transformer [33], a widely used attention-based framework for permutation-invariant set aggregation, and our proposed lightweight module equipped with irrep-specific attention. Both the *Vanilla Set Transformer* and SIMORGH further improved predictive performance over ensemble representations alone, indicating that explicitly learning relationships between conformations is more effective than treating them independently. On MISATO, both approaches achieved the highest overall performance (PR-AUC *≈* 0.53). On the more challenging PLINDER apo benchmark, they likewise outperformed both the static and ensemble-only variants, demonstrating that the benefits of learned ensemble integration generalize beyond holo-derived datasets.

Although the *Vanilla Set Transformer* achieves predictive performance comparable to SIMORGH, the proposed architecture provides a substantially lighter aggregation mechanism that is more specialized to the MOR input structure. By performing order-aware equivariant aggregation independently across irreps, SIMORGH uses fewer trainable parameters and can separately weight the signal carried by each irrep, rather than mixing them indiscriminately. Finally, sensitivity analysis showed that predictive performance increases rapidly as additional conformations are incorporated before saturating within the noise at approximately eight conformations (Fig. S3), suggesting that performance does not scale indefinitely with ensemble size.

### 2.3 SIMORGH remains competitive on conventional benchmarks while improving ligand-free prediction

To evaluate SIMORGH against state-of-the-art methods, we first used conventional holo benchmarks and then extended evaluation to a more challenging ligand-free setting. We first compared SIMORGH with R-EGNN [11], a dynamic framework that constructs heterogeneous edges from interatomic cosine similarity of dynamic correlations, on the MISATO dataset. Incorporating conformational dynamics improved both architectures over their respective static counterparts. Moreover, both SIMORGH Static and Dynamic achieved higher accuracy, precision, and F1 score than their corresponding R-EGNN model (Table 2).

**Table 2:** Performance comparison with R-EGNN over the MISATO dataset. Baseline results are taken from [11]; SIMORGH results represent the mean across three independent split seeds. Bold indicates the best performance.

| Model | Accuracy | Precision | Recall | F1 Score |
| --- | --- | --- | --- | --- |
| R-EGNN Static | 0.8038 | 0.2749 | 0.5368 | 0.3617 |
| R-EGNN Dynamic | 0.8387 | 0.3353 | **0.5498** | 0.4142 |
| SIMORGH Static | 0.9398 | 0.4289 | 0.4864 | 0.4194 |
| SIMORGH | **0.9454** | **0.4844** | 0.5028 | **0.4455** |

We next benchmarked SIMORGH against representative pocket-prediction methods, including P2Rank [34], DeepPocket [16], DeepSurf [35], and LABind [3] (Fig. 2). Performance was assessed in two complementary ways: as a binary classification task using PR-AUC, and as discrete binding-site prediction, obtained by applying HDBSCAN [36] clustering within a dedicated post-processing pipeline (Methods). For the latter, we used two geometric success metrics: DCA (distance from the predicted binding-site center to the nearest ligand atom) and DCC (distance between predicted and true binding-site centers), both with a threshold of 4 Å. On conventional holo benchmarks, SIMORGH achieved DCC performance comparable to existing methods, whereas on DCA, it predominantly outperformed them (Fig. 2a). The DCC improvements were less pronounced, as this point-to-centroid metric is inherently more sensitive to the rigid 4 ^Å^ cutoff when accommodating conformational variations across the ensemble.

**Figure 2:**
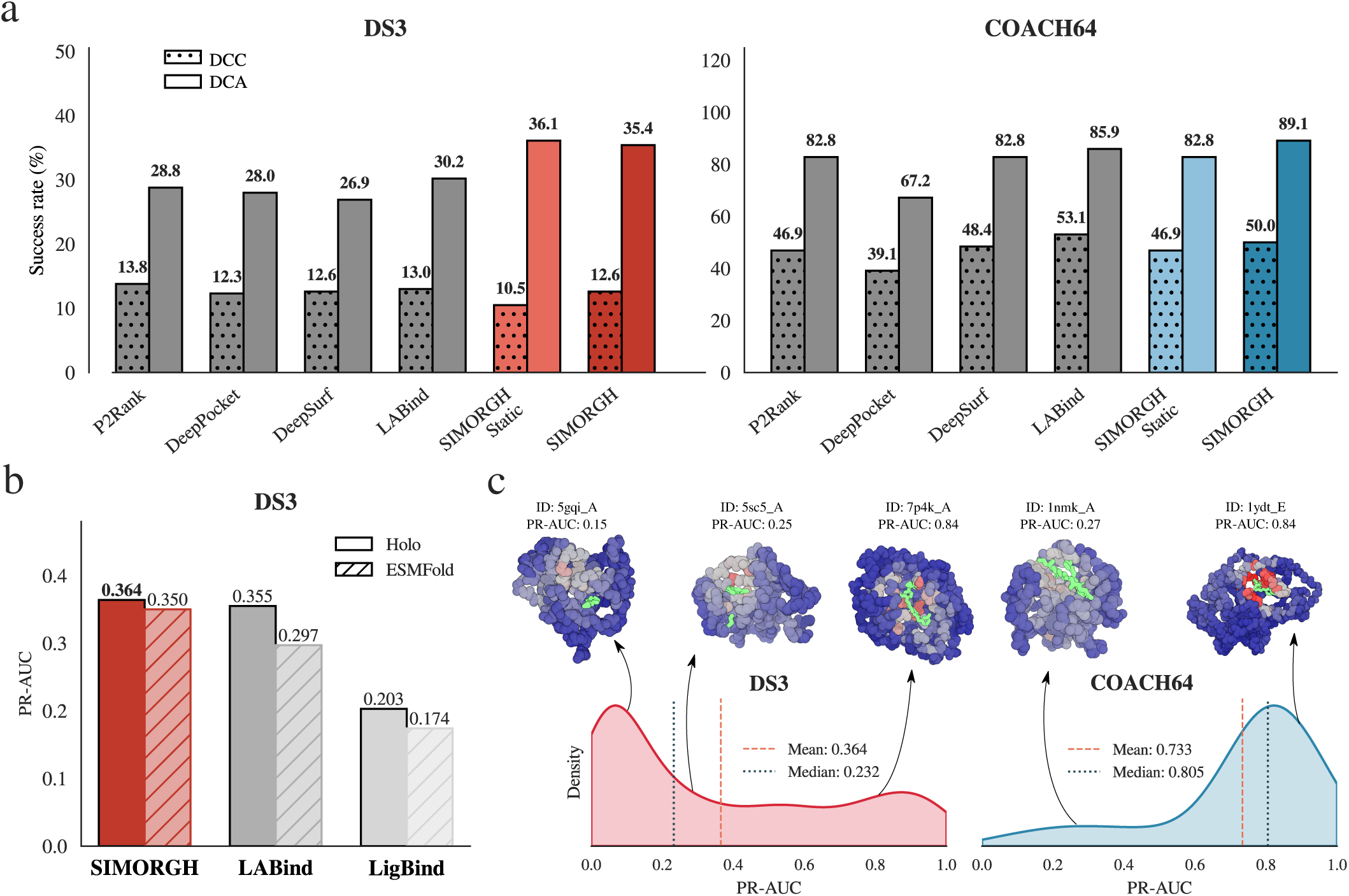
Comparison of SIMORGH with state-of-the-art ligand-binding-site prediction methods. **a**: Comparison with P2Rank, DeepPocket, DeepSurf, and LABind on the COACH64 and DS3 holo benchmarks using DCC and DCA success metrics. **b**: Performance on DS3 using experimentally determined holo and ESMFold-predicted structures. **c**: Representative successful predictions across varying structural complexities and challenging failure cases, with residue-level binding probabilities visualized on protein surfaces and PR-AUC reported for each example. Baseline values are from [3]; SIMORGH was evaluated on entries with successfully generated ensembles (Table S3).

Next, we hypothesized that in holo structures, conformational heterogeneity becomes partially redundant because ligand binding is already preorganized. This prevents SIMORGH from taking advantage of ensemble information. To further evaluate this, we considered a ligand-free setting using ESMFold-predicted structures that represent less-organized pockets and deviate further from the holo conformation [3]. In this scenario, a markedly different picture emerged. Here, the performance of existing methods dropped considerably, while SIMORGH retained nearly identical predictive accuracy between the holo and apo-like structures, outperforming both LABind and LigBind. (Fig. 2b).

Inspection of individual predictions supports this interpretation (Fig. 2c, Fig. S4): SIMORGH accurately localizes binding sites across proteins with diverse structural variability, while performance declines for multiligand systems and for exceptionally small ligands which have weak geometric signatures. Together, these results indicate that SIMORGH remains competitive on established holo benchmarks while generalizing more robustly to apo-like conditions, consistent with an ensemble-based learning strategy that captures conformational determinants of binding rather than relying on ligand-induced pocket geometry alone.

### 2.4 Ensemble-aware learning enables robust prediction of cryptic pockets from apo proteins

The ultimate objective of ligand binding-site prediction is to identify functional pockets directly from ligand-free proteins. This remains particularly challenging for cryptic pockets, which are absent or only weakly formed in experimentally resolved apo structures and emerge only after ligand binding or through transient conformational fluctuations. To determine whether ensemble-aware learning overcomes this limitation, we evaluated SIMORGH on CryptoBench, a recently introduced benchmark comprising experimentally resolved apo–holo pairs and cryptic binding sites [37]. SIMORGH was trained using BBFlow-derived conformational ensembles while maintaining a strict sequence identity threshold (*<* 30%) between training and test proteins.

To disentangle the contribution of ensemble type from model quality, we employed four input modalities: BBFlow-derived ensembles of holo structures, BBFlow-derived ensembles of apo structures, MD simulation trajectories of apo structures, and AlphaFlow-generated ensembles of single-chain entries, the latter motivated by recent evidence that static AlphaFold structures approximate the apo conformational state [38]. We then assessed the structural bias of each ensemble generation method by comparing them to native apo and holo structures (Fig. S5, S6). As expected, BBFlow ensembles remained strongly biased toward their input state, with apo- and holo-derived ensembles predominantly resembling their respective starting conformations. MD trajectories sampled a broader conformational space (Fig. S7) and deviated more substantially from the apo input structure, while retaining an overall apo preference. AlphaFlow ensembles, in contrast, showed no clear bias toward either state, exhibiting comparable similarity to native apo and holo conformations and suggesting more balanced sampling of the apo–holo transition landscape.

We therefore benchmarked SIMORGH directly against P2Rank and PocketMiner[39] on single-chain apo/holo pairs across all four input configurations (Fig. 3a). Because P2Rank and PocketMiner require full explicit atomic coordinates, they were evaluated on the corresponding native apo or holo structures when compared with SIMORGH using BBFlow-generated backbone-only ensembles, while for the AlphaFlow and MD configurations they were evaluated as mean performance across their respective ensembles. As expected, all models performed best on holo structures, with SIMORGH and P2Rank achieving comparable top-tier performance (PR-AUC of 0.52 and 0.51, respectively). Performance declined for all models on unbound structures, but the decay was substantially smaller for SIMORGH, which maintained a PR-AUC of 0.46 on BBFlow-generated ensembles derived from native apo structures, compared to 0.37 for P2Rank evaluated on the corresponding single native apo structures. Notably, MD trajectories yielded performance on par with native apo structures across all models, without any benefit from naive ensemble averaging for the standard predictors, underscoring that SIMORGH’s gains stem from learned heterogeneity integration rather than broader input coverage, and that this advantage generalizes to physically realistic trajectories despite the model being trained exclusively on BBFlow-derived data. AlphaFlow ensembles produced the lowest performance across all models, consistent with their conformational profile lying furthest from any single reference state; yet SIMORGH retained the highest accuracy in this setting as well (PR-AUC of 0.39). Collectively, these results establish that SIMORGH is better adapted than existing methods to the ligand-absent conditions that characterize realistic drug discovery workflows.

**Figure 3:**
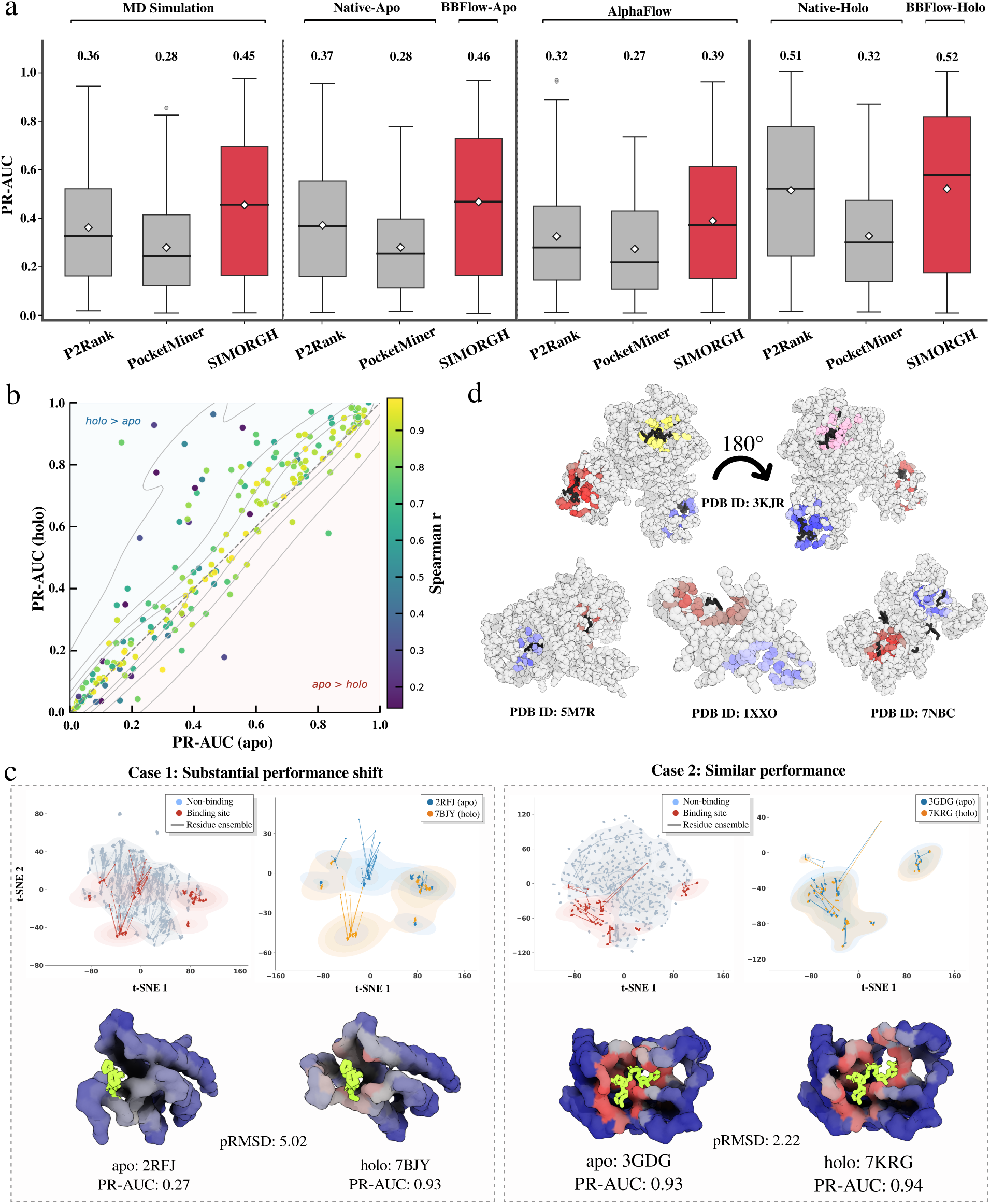
**a**: Performance comparison between P2Rank, PocketMiner, and SIMORGH based on single-chain PR-AUC distributions with mean values. Sample sizes are: AlphaFlow (n=174 for all); native structures (n=174 for SIMORGH/P2Rank; n=155 and n=165 for PocketMiner apo and holo); BBFlow ensembles (n=174); and MD simulation clusters (n=172 for all). **b**: Predictive consistency across BBFlow-derived apo and holo ensembles after outlier removal. Points represent individual targets, colored by the Spearman correlation (r) computed over the union of apo and holo binding-site residues. **c**: Conformational determinants of latent space stability for Case 1 (PDB 2RFJ/7BJY) and Case 2 (PDB 3GDG/7KRG). Top: Residue-level t-SNE embeddings where each node represents a single residue in one conformation, and identical residues are linked across conformations via lines (blue: non-binding; red: binding site). Bottom: Corresponding pocket environments colored by prediction confidence, with co-crystallized ligands in green. **d**: Visualized structural examples of multi-chain and interface pockets (PDB IDs: 3KJR, 5M7R, 1XXO, 7NBC).

Given these results, we next asked whether this advantage reflects genuinely consistent, conformation-aware predictions across apo and holo ensembles of the same target. SIMORGH exhibited strong predictive consistency, with a global residue-level Spearman correlation of 0.87 across all residues, remaining substantial (0.73) when restricted to binding-site residues (Fig. 3b). This consistency was not merely an artifact of structural rigidity: predictive agreement correlated only weakly with pocket root-mean-square deviation (pRMSD) between apo and holo states (Spearman *ρ* = *−*0.32; Fig. S8), confirming that robust predictions are maintained even for conformationally flexible targets.

To understand the mechanistic basis for this consistency, and for the cases where it breaks down, we analyzed residue-level t-SNE embeddings alongside structural data for two contrasting targets: one where apo performance degrades sharply relative to the holo state (Case 1), and one where high accuracy is preserved across both conformations (Case 2) (Fig. 3c). In Case 1, the model fails to linearly segregate binding from non-binding residues in latent space, and the apo and holo representations of the binding site diverge rather than align. This target undergoes a large structural rearrangement upon ligand binding (pRMSD of 5.02 Å, versus 2.22 ^Å^ for Case 2), which perturbs the local geometry enough to cause misclassification. In Case 2, by contrast, pocket topology is preserved across the conformations sampled, and the model maintains a stable, conformation-invariant embedding space that supports accurate classification. Together, these cases indicate that SIMORGH succeeds when the sampled conformational ensemble adequately covers the neighborhood of the binding-competent state, allowing it to average away the noise.

Beyond single-chain targets, multimeric proteins present a distinct challenge, as functional pockets can be buried within oligomeric interfaces, compounding the difficulties of size and complexity inherent to multi-chain assemblies. SIMORGH successfully identified both pockets localized within individual subunits and those spanning multiple chain interfaces (Fig. 3d). Nonetheless, multimeric targets showed a marked performance reduction, with mean PR-AUC falling from 0.46 for single-chain apo entries to 0.36 for multi-chain entries. This decline reflects two contributing factors: geometrically valid, high-confidence pocket predictions that lack a co-crystallized ligand in the reference structure, and shallow surface clefts that occasionally escape detection owing to the absence of deeply buried geometric signatures.

### 2.5 Integration with Galaxy enables accessible and reproducible deployment

To facilitate broad adoption and comply with FAIR (Findable, Accessible, Interoperable, and Reusable) principles, SIMORGH has been integrated into the Galaxy toolshed [40] and deployed on the European Galaxy server (https://usegalaxy.eu/) [41]. The complete workflow, from protein structure input to ligand binding-site prediction, is accessible through a web-based interface without requiring local software installation, dedicated computational infrastructure, or command-line expertise. Optionally, users can generate backbone conformational ensembles using BBFlow, perform HDBSCAN-based binding-site clustering, and interactively visualize predicted binding pockets within the same workflow. To ensure portability and reproducibility across computing environments, SIMORGH is distributed as a Docker container [42] and published through the Galaxy ToolShed, enabling straightforward deployment on local, institutional, or cloud-based Galaxy instances. A step-by-step training was also created in the Galaxy training platform [43], where users can go through a dedicated tutorial available at https://gxy.io/GTN:T00594 and learn how to use SIMORGH and interpret the results.

## 3 Discussion

Protein–ligand binding site prediction has long been constrained by two compounding limitations: reliance on single static conformations and evaluation on holo structures whose pocket geometry is already pre-organized around the ligand. Our results suggest that these limitations stem from a common underlying assumption that ligand binding can be inferred solely from an individual protein structure. Instead, the consistent improvements that we observed across realistic apo-state evaluations and cryptic pocket prediction indicate that ligand binding can be better viewed as a property of a protein’s conformational ensemble. By considering conformational heterogeneity as a source of information, SIMORGH demonstrates that integrating complementary geometric evidence across multiple conformations substantially improves generalization under the ligand-free conditions encountered in current drug discovery pipelines.

A central aspect of SIMORGH is the explicit separation of geometric representation learning from ensemble reasoning. This two-stage formulation allows conformational information to contribute in two complementary ways. First, learning from multiple conformations naturally augments the geometric encoder, exposing it to diverse forms of the same protein and yielding transferable MORs that are less dependent on any individual structure. Second, the ensemble aggregation stage explicitly reasons across these representations to distinguish persistent structural signals from transient conformational fluctuations. The resulting performance gains consistently exceeded those obtained through increased architectural complexity alone, suggesting that the principal limitation of existing binding-site predictors is not insufficient model capacity, but insufficient structural information.

The source and scale of conformational sampling also influence model performance. MD-derived ensembles consistently provide richer training signals than alternative generative models, as they can better capture the physical conformational landscape relevant for ligand recognition. Nevertheless, the final model, trained exclusively on BBFlow-generated backbone ensembles, generalizes robustly to MD trajectories during inference. This observation suggests that SIMORGH learns to exploit geometric heterogeneity as an intrinsic structural signal rather than overfitting to the statistical properties of a particular sampling strategy. At the same time, MD simulation remains the most physically realistic representation of protein dynamics, and training on comparably large MD-derived datasets remains an important direction for future work, one that is becoming increasingly feasible through the rapid expansion of public MD repositories [5–8].

Despite these advances, an important limitation reflects a broader challenge for the field: the absence of well-defined ground truth for the ligand-free binding landscape. Current benchmark datasets generally annotate residues observed to contact ligands in one or a small number of experimentally resolved structures, whereas proteins populate continuous conformational ensembles whose binding competence varies over time. Consequently, all supervised learning approaches, including SIMORGH, are necessarily trained against an incomplete representation of the underlying functional landscape. By learning directly from conformational ensembles, SIMORGH partially mitigates this limitation by reducing the tendency to learn ligand-induced structural rearrangements from individual holo structures and instead modelling the distribution of conformational states surrounding ligand binding. Nevertheless, our latent-space analysis demonstrates that individual conformations differ substantially in their informativeness, with some conformations carrying strong predictive signals while others primarily represent thermal fluctuations or structurally ambiguous intermediate states. Constructing dedicated apo-state datasets in which binding annotations are systematically transferred across experimentally validated conformational ensembles would therefore provide a substantially stronger foundation for future ensemble-aware learning methods.

Beyond ligand binding-site prediction, our results establish a broader design principle for geometric machine learning over structurally heterogeneous biological systems. The consistent superiority of explicit ensemble reasoning over other models indicates that structural variability should be treated as a first-class learning signal rather than as an auxiliary feature. Although demonstrated here for ligand binding-site prediction, the same principle naturally extends to intrinsically disordered proteins, allosteric regulation, protein–protein interaction interfaces, conformational state classification, and other problems in which biological function emerges from distributions of structures rather than a single conformation [2]. As increasingly large collections of experimentally determined and computationally generated conformational ensembles become available, scalable ensemble-aware equivariant architectures such as SIMORGH provide a practical foundation for next-generation geometric learning across proteome-scale structural datasets.

## 3.1 Data and code availability

All model variants, benchmarking results, and statistical analyses are available on GitLab at https://gitlab.inria.fr/delta/SIMORGH. SIMORGH can be run directly using a Google Colab notebook (https://colab.research.google.com/drive/1FSkESlIdVksg2d0eBlj0E1AmxI9Hc1bc) or the Docker container https://ghcr.io/amisteromid/simorgh:v1.0.0. It is also available through the Galaxy ToolShed https://toolshed.g2.bx.psu.edu/view/bgruening/simorgh/673894ae03a8 and can be executed on the public Galaxy server at https://usegalaxy.eu/?tool_id=simorgh. The data are also available on Zenodo at https://doi.org/10.5281/zenodo.18414049

## 4 Author Contributions

OM, YK, and HK conceived the initial idea. OM developed the SIMORGH architecture in discussion with YK and HK, and performed benchmarking and the data analyses. HK and YK supervised the project. ANN created the Docker container and integrated SIMORGH into Galaxy with supervision from BG. OM and HK wrote the initial draft. All authors contributed to writing the manuscript.

## 5 Acknowledgements

HK was supported by the French National Research Agency (ANR), under grants ANR-22-CPJ2-0075-01 and ANR-24-CE45-4243-01. YK was supported by the French National Research Agency (ANR) under the France 2030 grant reference number ANR-24-RRII-0002, operated by the Inria Quadrant Program. ANN and BG were funded by the German Federal Ministry of Research, Technology, and Space BMFTR grant 031 A538A de.NBI-RBC and the Ministry of Science, Research and the Arts Baden-Württemberg (MWK) within the framework of LIBIS/de.NBI Freiburg. This work was granted access to the HPC resources of IDRIS under the allocation 2024-A0160715132, 2025-A0180714660 and 2026-A0200714660 made by GENCI.

## 7 Methods

### 7.1 Dataset and ground truth annotation

Model optimization, primary evaluation, and comparison with the dynamic-aware R-EGNN were performed using MD trajectories from the MISATO database [6]. Protein conformations were first structurally aligned using the Biopython structural superimposer [44]. Complexes lacking ligands, or containing ligands that exhibited substantial translational drift during simulation (indicative of an unstable binding mode) were excluded from all analyses. Binding-site residues were defined by a distance cutoff of 4.5 ^Å^ between protein and ligand heavy atoms, and ground-truth annotations were derived from the first frame of each MD trajectory as a representative of the equilibrium binding state.

To ensure strict separation between training, validation, and test sets, dataset redundancy was removed via a two-step sequence clustering procedure using MMseqs2 [45]. Protein-ligand complexes were first filtered at 90% sequence identity, yielding 3,456 non-redundant entries. The remaining proteins were then clustered at 30% sequence identity, and entire clusters were assigned to either the training, validation, or test partition, preventing any homologous proteins from appearing across splits. To assess the stability of reported metrics, three independent models were trained using different random seeds and dataset partitions. To validate the optimized representation order on single-chain proteins independently of the Misato setting, additional experiments were conducted using the PUResNetV2.0 training dataset [18], for which conformational ensembles were generated with AlphaFlow [15]. After redundancy reduction at 30% sequence identity, this dataset comprised approximately 8,000 single-chain protein structures.

To scale training beyond the limited availability of MD trajectories, we incorporated backbone conformational ensembles generated by BBFlow [14], a state-of-the-art generative ensemble predictor applicable to both single-chain proteins and multi-chain complexes. Structures exceeding 1,900 residues were excluded due to memory constraints. Non-standard amino acids and residues with missing backbone atoms were removed before processing, and all conformations were realigned to a common reference frame. We constructed two benchmark suites to enable direct and fair comparison with prior works (Supplementary Figure S9). First, following Zhang et al. [3], the DS3 dataset was used to evaluate binding-site prediction under strict protein-ligand separation, with an additional subset of COACH64 used for evaluation. Second, the Crypto-Bench dataset [37] served as the benchmark for cryptic pocket prediction. For this setting, the training set combined DS3 with several datasets commonly used in binding-site prediction studies, namely scPDB [46], COACH420, and HOLO4K [34], supplemented with BBFlow-derived ensembles of the CryptoBench training entries (Table S3). To prevent information leakage, any training protein sharing more than 30% sequence identity with a CryptoBench test protein was excluded. BBFlow was used to generate backbone ensembles for both the apo and holo structures of each CryptoBench protein pair. For the test set specifically, two additional sources of conformational input were generated to enable cross-modality evaluation: AlphaFlow-derived ensembles and all-atom MD simulation trajectories. MD simulations were performed on apo-state structures using three independent replicas of 500 ns each, providing a physically grounded reference ensemble for assessing cross-protocol generalization. Table S4 summarizes the number of conformations generated per method throughout this work.

Ground-truth labels for BBFlow-derived structures were generated using an automated binding-site annotation pipeline that integrates structural information from PDB coordinate files with biologically curated ligand annotations from Q-BioLiP [47]. Only protein chains associated with biologically relevant ligands were retained. Ligand identity was established by first matching the chemical component annotations provided by Q-BioLiP; when no matching ligand entry could be identified, the ligand chain geometrically closest to the receptor protein was selected as a fallback. Binding-site residues were then defined as those containing at least one heavy atom within 4.5 ^Å^ of the identified ligand, and these annotations were mapped onto the protein sequence to generate binary residue-level classification labels.

### 7.2 Molecular dynamics simulations setup

Three independent 500-ns MD replicates were performed for each of the 207 apo-state structures from CryptoBench, starting from the corresponding experimental structures. Missing residues were modeled using MODELLER [48]. Simulations were run with GROMACS 2024.2 [49] and the CHARMM36m force field. Each system was placed in a dodecahedral box of explicit TIP3P water with a minimum buffer distance of 12 ^Å^ from the solute, and Na^+^ and Cl*^−^* ions were added to reach physiological salt concentration (150 mM NaCl). Hydrogen atoms were added and histidine protonation states assigned using Reduce [50]. Before production runs, each system underwent energy minimization by steepest descent for 5000 steps, followed by NPT equilibration at 310 K, during which harmonic positional restraints on all protein and lipid non-hydrogen atoms were progressively released over six steps, totaling 0.375 ns. Pressure was controlled at 1 bar using the Berendsen barostat during equilibration [51]. Production simulations were carried out in the NPT ensemble with a 2 fs time step, with three replicates per system generated using different initial velocities. Temperature was maintained at 310 K with the V-rescale thermostat [52] and pressure at 1 atm with the C-rescale barostat under isotropic conditions [53]. Bonds involving hydrogen atoms were constrained with LINCS [54], long-range electrostatics treated with Particle Mesh Ewald [55], and coordinates saved every 100 ps. In total, this protocol yielded 310.5 ms of aggregate MD simulation time, sampling more than 3 million conformations across all systems.

### 7.3 Evaluation metrics

To enable comparison with previous studies, two complementary evaluation metrics are reported.

1. Pocket-centric evaluation including two geometric criteria: DCC (distance between predicted and true binding-site centers) and DCA (distance between the predicted pocket center and the closest ligand atom). For each metric, a prediction is considered successful if the distance is below 4 Å.
2. Residue-based evaluation including standard classification metrics: precision, recall, F1 score, MCC, and PR-AUC, computed directly from predicted residue-level probabilities. Given the strong class imbalance and the relative nature of binding-site detection, PR-AUC is used as the primary performance metric, as it better reflects ranking quality between binding-site and non-binding-site residues than threshold-dependent metrics.

### 7.4 Features

We encode the static protein structures using a focused set of node and edge features. We construct a residue-level graph where nodes correspond to residues and edges are defined between residues whose C*α* atoms are within an 11 Å distance threshold. Let **x***_i_ ∈* R^3^ denote the C*α* coordinate of residue *i*, and let 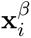 denote the corresponding C*β* coordinate.

#### Node features

Node features consist of the residue identity (type) and the side-chain orientation vector (SCOV). For each residue, the SCOV is defined as:

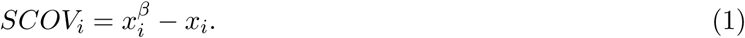

To ensure consistency for glycine, which lacks a C*β* atom, a pseudo-C*β* position is constructed for all residues using a planar backbone approximation. Given backbone atoms N, C*α*, and C, the orientation direction is defined as:

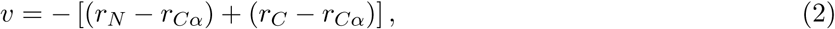

and the pseudo C*β* is placed along the normalized direction **v**^ at a bond length of 1.52 Å.

#### Edge features

Edge features capture the spatial relationship between connected nodes and include the relative displacement vector (*R*), the pairwise C*α* distance (*D*), and the side-chain orientation difference (SCOD).

For residue *i* and its neighbor *j ∈ N* (*i*), we define the relative displacement vector *R* as:

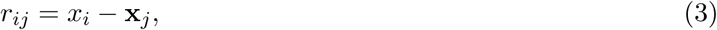

and the corresponding Euclidean distance *D* as:

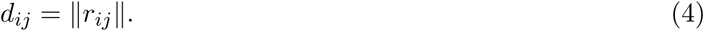

To characterize the relative side-chain packing of residues, we define the side-chain orientation difference (SCOD) as:

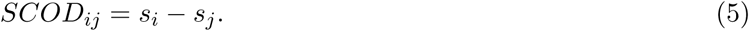

### 7.5 Architecture

Here, we describe the SIMORGH architecture by first introducing the core equivariant operators and representation layers, followed by a step-by-step description of the data flow across the encoder, message construction, attention-based aggregation, decoder, and ensemble aggregation modules, and finally the training and inference procedure.

#### Spherical Harmonic Encoding

For vectorial features, we employ spherical harmonics *Y_ℓm_*(^**r**) to map spatial information into the irrep basis. As the eigenfunctions of the Laplacian on the sphere *S*^2^, they form the canonical equivariant basis for angular functions. The projection *Y* ^(^*^ℓ^*^)^ : *S*^2^ *→* R^2^*^ℓ^*^+1^ encodes the orientation of relative position vectors ^**r***_ij_*, allowing the network to capture anisotropic interactions and higher-order multipoles while strictly maintaining equivariance.

#### Bessel Basis Layer

To represent interatomic distances, we adopt a radial basis construction inspired by GemNet [56], using a Bessel basis with a smooth polynomial envelope (Supplementary Figure S10). Specifically, distances are expanded using sinusoidal Bessel functions with learnable frequencies, modulated by an envelope function that enforces a smooth cutoff at the interaction radius. The envelope ensures that both the basis and its derivatives vanish continuously at the cutoff to improve numerical stability. The resulting radial features are further processed by a lightweight Multilayer Perceptron (MLP) with optional layer normalization and offset, enabling flexible yet well-regularized distance-dependent message modulation.

#### Depth-wise Tensor Product

Inspired by [30, 57], we implement a depth-wise tensor product by constructing instructions for all valid Clebsch-Gordan couplings *l*_in_ *⊗ l*_edge_ *→ l*_out_ required by the target output irreps. As a result, each output vector at a given multiplicity index depends exclusively on the corresponding vectors from each input irrep feature through valid Clebsch–Gordan couplings, illustrating the multiplicity-preserving (depth-wise) nature of the tensor product.

#### Equivariant Linear Layer

As in Equiformer [30], linear layers act independently on each irrep type *L*, with biases only for scalars (*L* = 0) to preserve equivariance.

#### Equivariant Layer Normalization

We apply an equivariant layer normalization over irreps, following Equiformer [30]. Input is decomposed into irrep blocks and each block is normalized independently. For each irrep block, scalar features are first centered by subtracting their mean across channels. Then, for all irreps, a normalization factor is computed from the average squared magnitude of the vectors within the block, and the block is rescaled accordingly. Learnable scale parameters are applied per channel, and for scalar features an additional learnable bias is used.

The network processes these primitives through the following modules:

#### Encoder

The encoder initializes node representations by concatenating chemical semantics (residue identity) with explicit side-chain orientation vectors (SCOV). To lift these inputs into node-level MOR, we apply a self-interacting fully connected tensor product (FCTP) that acts as a geometric expansion layer. To incorporate local geometric context, the encoder then aggregates neighborhood information through a depthwise tensor product operating on neighboring node embeddings and relative side-chain orientation differences (SCOD), while radial distances modulate interaction strength through learnable radial basis functions. The aggregated neighborhood features are added to the self-interaction embedding to form the initial node-level MOR.

#### Raw Message

To construct the raw messages, we explicitly decompose each neighbor’s hidden state into irrep-specific subspaces. Scalar features (*ℓ* = 0), which primarily encode stable chemical semantics, are propagated directly and form the scalar raw message (SRM). Directional features (*ℓ* = 1) require geometric alignment and are therefore processed through a depthwise tensor product. Specifically, the source node’s vector features interact with edge-level geometric descriptors, consisting of the relative displacement vector **r***_ij_* and SCOD*_ij_*. This interaction is modulated by a learnable radial function of the inter-residue distance, producing a vector-derived raw message (VRM) that captures the spatially and orientationally weighted influence of the neighbor. The resulting tensor product lifts vector features into higher-order components according to the Clebsch–Gordan decomposition 1 *⊗* 1 *→* 0 *⊕* 1 *⊕* 2.

#### Equivariant Graph Transformer Aggregation

We introduce an *SE*(3)-equivariant attention-based aggregation mechanism for graph transformers. Queries (*Q*) and keys (*K*) are constructed as MOR via linear equivariant projections of node states and raw messages into matched representation spaces. Multi-head structure is obtained by projecting features into a direct sum of *H* head spaces. For each attention head *h*, attention coefficients are computed via an inner product

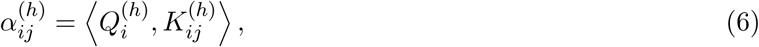

which yields a scalar by construction. This invariance permits the use of standard softmax normalization over each node’s neighborhood.

Value tensors *V_ij_* are computed using a receiver-aware transformation module. Instead of propagating sender messages directly, this module conditions message updates on the destination node state. The module consists of two components:

1. Decoupled Tensor Interactions: Scalar (*ℓ* = 0) and vector (*ℓ* = 1) components of the receiver node state are processed independently via depthwise tensor products, each interacting exclusively with the corresponding components of the raw message to have order-specific refinement of geometric features.
2. Equivariant Gating Nonlinearity: The resulting tensors are concatenated and passed through a gated equivariant nonlinearity, where learned scalar channels modulate higher-order components.

The gated value tensors are projected back into the node MOR space, distributed across attention heads, and weighted by the invariant attention coefficients. Aggregation is performed via scatter summation over incoming edges, followed by head recombination. A residual connection adds the aggregated message to the original node state, completing the equivariant update.

#### Decoder

To translate MORs into binding-site probabilities, we use a two-step decoder. First, MORs representations are processed using a self-interacting depthwise tensor product constrained to produce only scalar (*ℓ* = 0) channels. The resulting scalar features are then passed through a standard MLP with SiLU activations to produce the final binding-site likelihood scores.

#### MOR Set Aggregation

For residue *i*, let 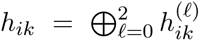 its MOR in conformation *k ∈* {1*, … , K*}. We first construct an initial consensus anchor by mean-field aggregation,

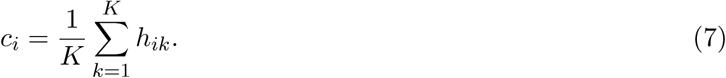

Each conformation is then compared with the corresponding consensus anchor through a tensor-product-based geometric querying operation, producing an invariant compatibility score for each tensor order,

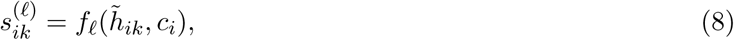

where *h̃_ik_* denotes the projected and gated representation. The scores are normalized across conformations independently for each residue and tensor order,

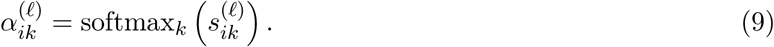

The resulting weights are applied to the corresponding irreducible components to obtain the refined ensemble representation,

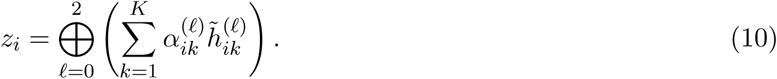

Finally, the refined representations are mean-field aggregated to obtain the global ensemble descriptor, which is normalized and passed through the decoding module to produce the final scalar output.

Training details are as follows:

#### Implementation

SIMORGH is implemented using PyTorch and the e3nn framework. At the framework level, equivariant linear and bilinear operations are implemented using tensor product modules adapted from Equiformer [30]. These layers are based on e3nn.o3.TensorProduct but modify the normalization, initialization, and bias handling to improve training stability. The hyper-parameters for base message passing model and set aggaretion model is summarized in Table S5.

#### Optimiser and Loss

We use a linear warm-up followed by cosine annealing [58] for the learning rate, with minimum value *η*_min_, and the focal loss [59] with tuned *α* and *γ* to mitigate class imbalance.

### 7.6 Clustering for Task Conversion

To convert residue-level binding predictions into binding-site coordinates, we applied HDBSCAN, a hierarchical density-based clustering algorithm that identifies stable clusters across multiple density levels [36, 60].

In contrast to standard HDBSCAN implementations that use only spatial proximity, we incorporated the model’s per-residue binding probabilities to bias clustering toward high-confidence predictions. We applied a lenient probability threshold of 0.4 while continuously penalizing lower-confidence residues. When all residues are classified as noise, a fallback binding-site center was defined as the geometric centroid of all residues with predicted probabilities exceeding 0.4. We modify the pairwise distance matrix as:

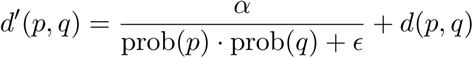

where (*d*(*p, q*)) denotes the Euclidean distance between residue coordinates, *prob*(*p*) and *prob*(*q*) are the predicted binding probabilities, *α* controls the contribution of the probability-dependent penalty term, and *ɛ* is a small constant added for numerical stability. *α*, together with the HDBSCAN hyperparameters, was optimized via a grid search over 294 parameter combinations (Fig. S11).

## 8 Supplementary Materials

## 8.1 Supplementary Figures

**Figure S1:**
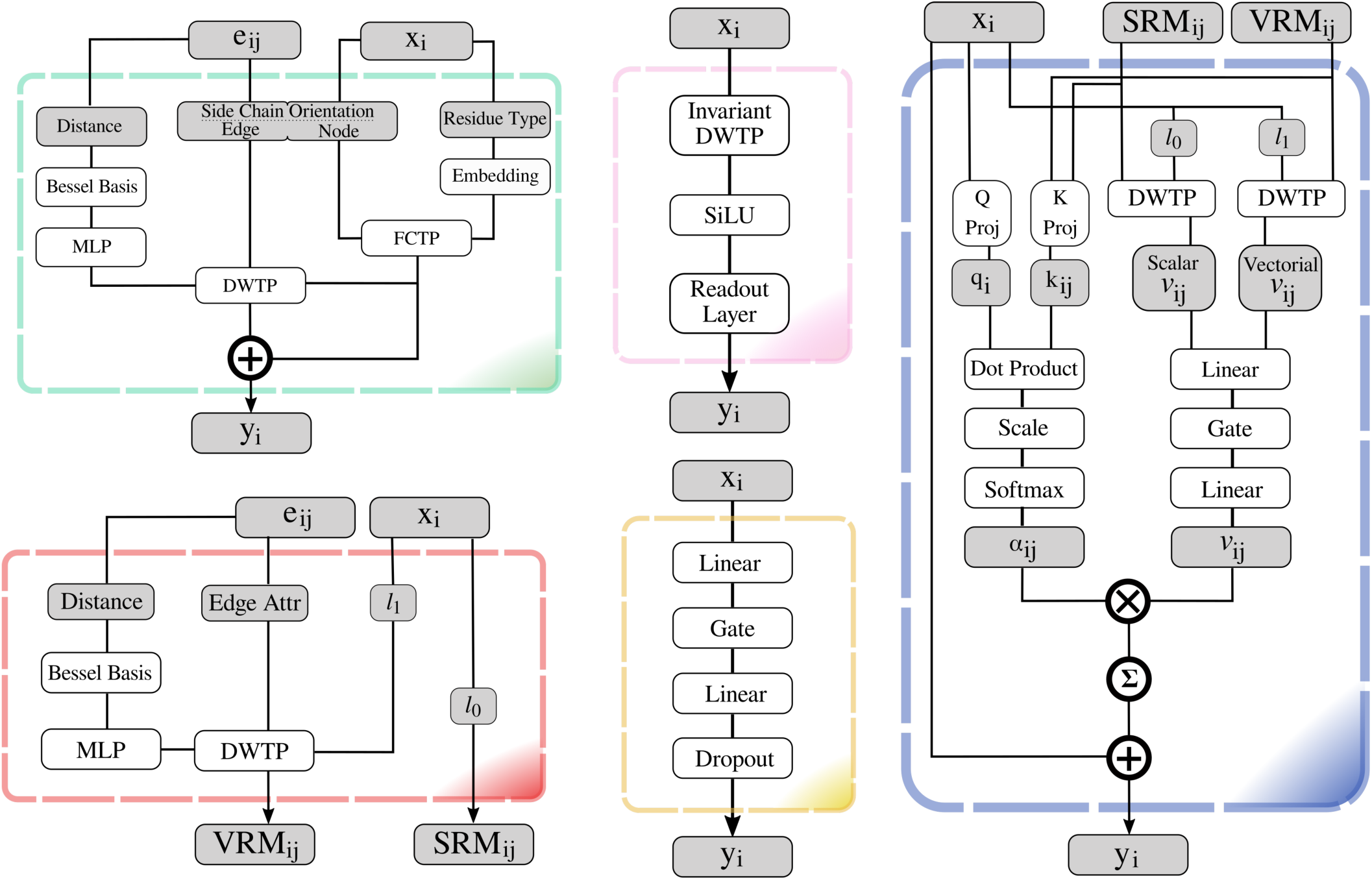
Notation used in the architecture diagrams: “*⊗*” denotes multiplication, “*⊕*” addition, and (in circle) neighbor aggregation. DWTP and FCTP refer to depth-wise and fully connected tensor products. SRM and VRM denote scalar and vector-derived raw messages, respectively. Block colors correspond to those in Figure 1a.

**Figure S2:**
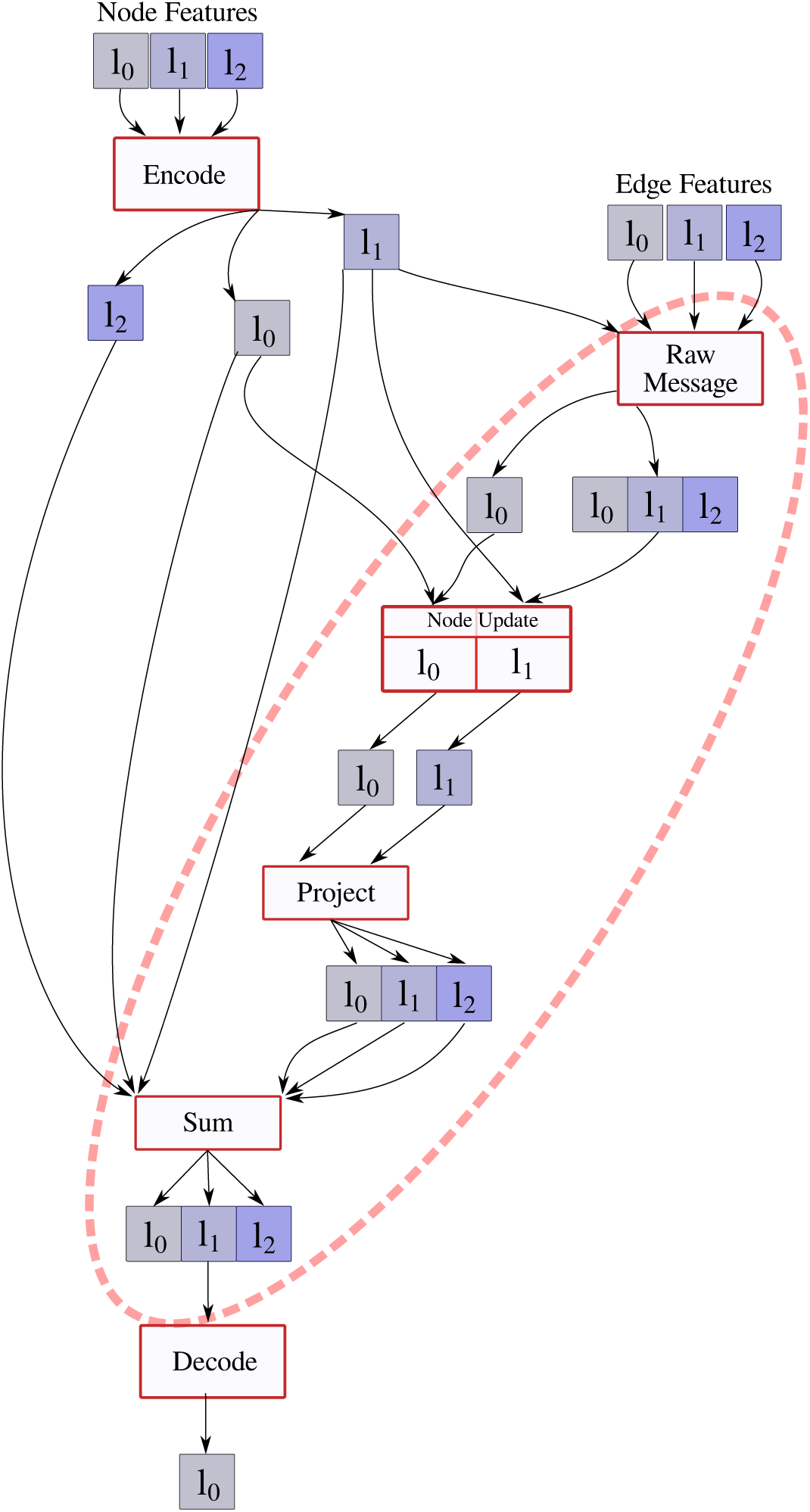
Node and edge features (*l*_0_*, l*_1_*, l*_2_) are embedded as MORs using spherical harmonics. Each block is defined by explicit input and output irreps, corresponding to the allowed coupling rules under the Clebsch–Gordan decomposition. In practice, not all theoretically permitted output irreps are instantiated in every module; for example, during the node update (message aggregation), vector-derived messages contribute only to the *l*_1_ channel, while the *l*_0_ and *l*_2_ channels are intentionally not computed. The dashed red loop highlights the iterative message-passing cycle, and finally, the resulting representations (*l*_0_*, l*_1_*, l*_2_) are then passed to the Decode module to produce per-residue predictions in the scalar *l*_0_ channel. This design reduces the number of tensor-product pathways by enforcing channel-selective cross-order interactions rather than fully coupling all irreps at every stage.

**Figure S3:**
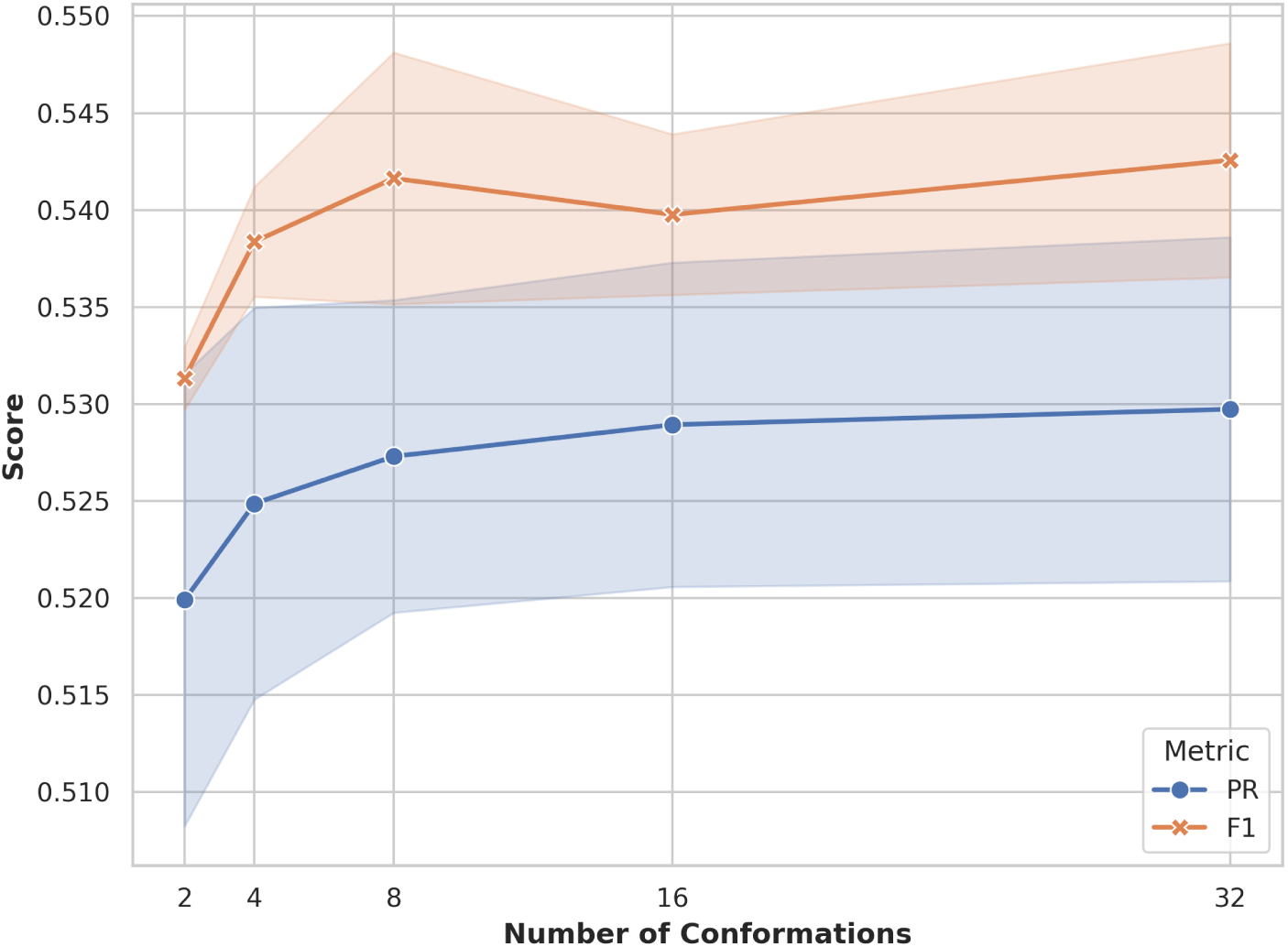
Model performance evaluated using PR-AUC and *F*_1_ score as a function of the number of input conformations (*K* = 2, 4, 8, 16, 32) included in the ensemble. The shaded regions denote the corresponding standard error (SE). This sensitivity analysis was conducted on the subset of the MISATO test set (*n* = 100 protein entries) to determine the optimal trade-off between the number of input conformations and predictive accuracy.

**Figure S4:**
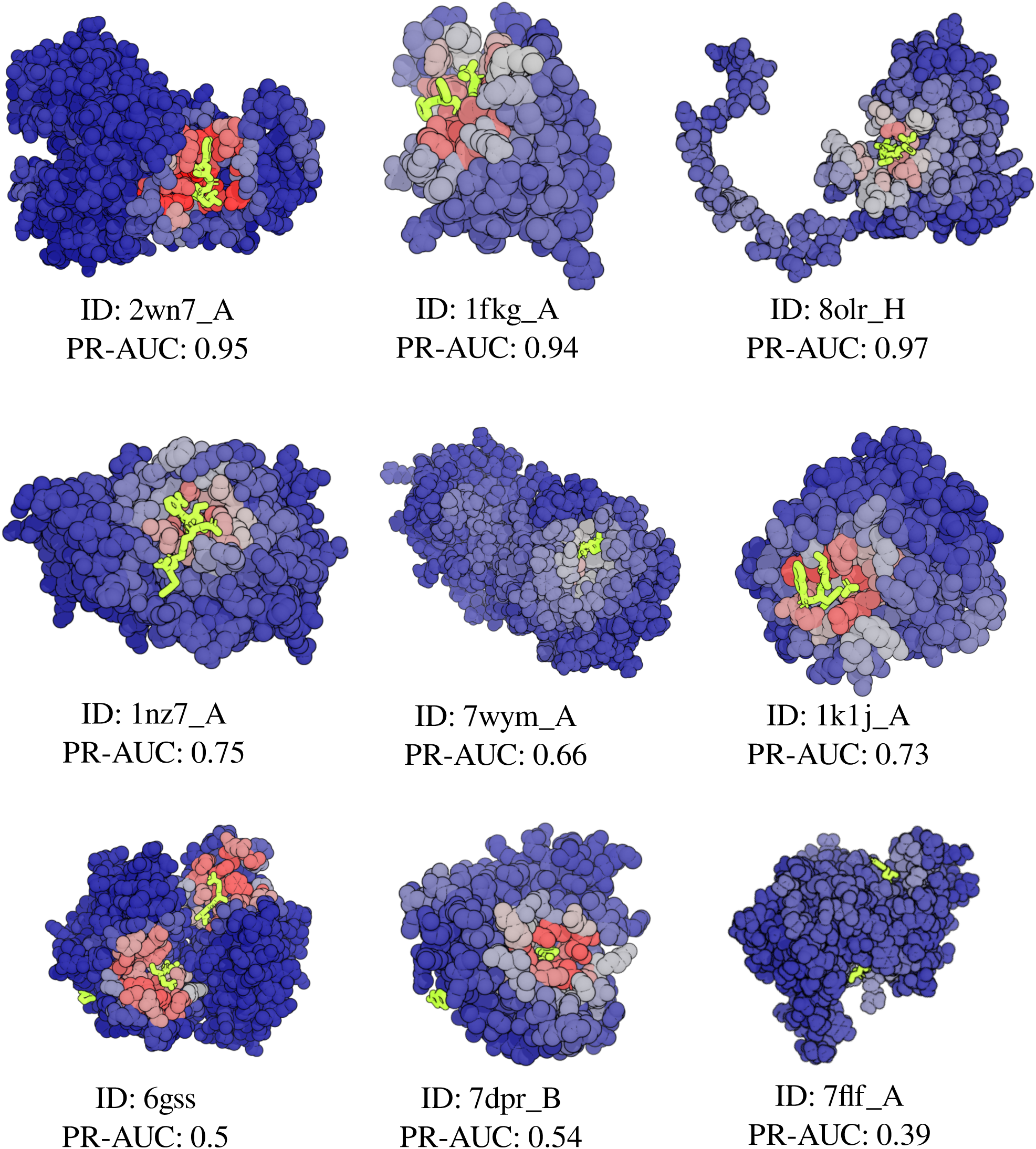
Visualization of model predictions, with a blue-to-red spectrum showing predicted probabilities. The ligand is shown in green, and the PR-AUC score is reported.

**Figure S5:**
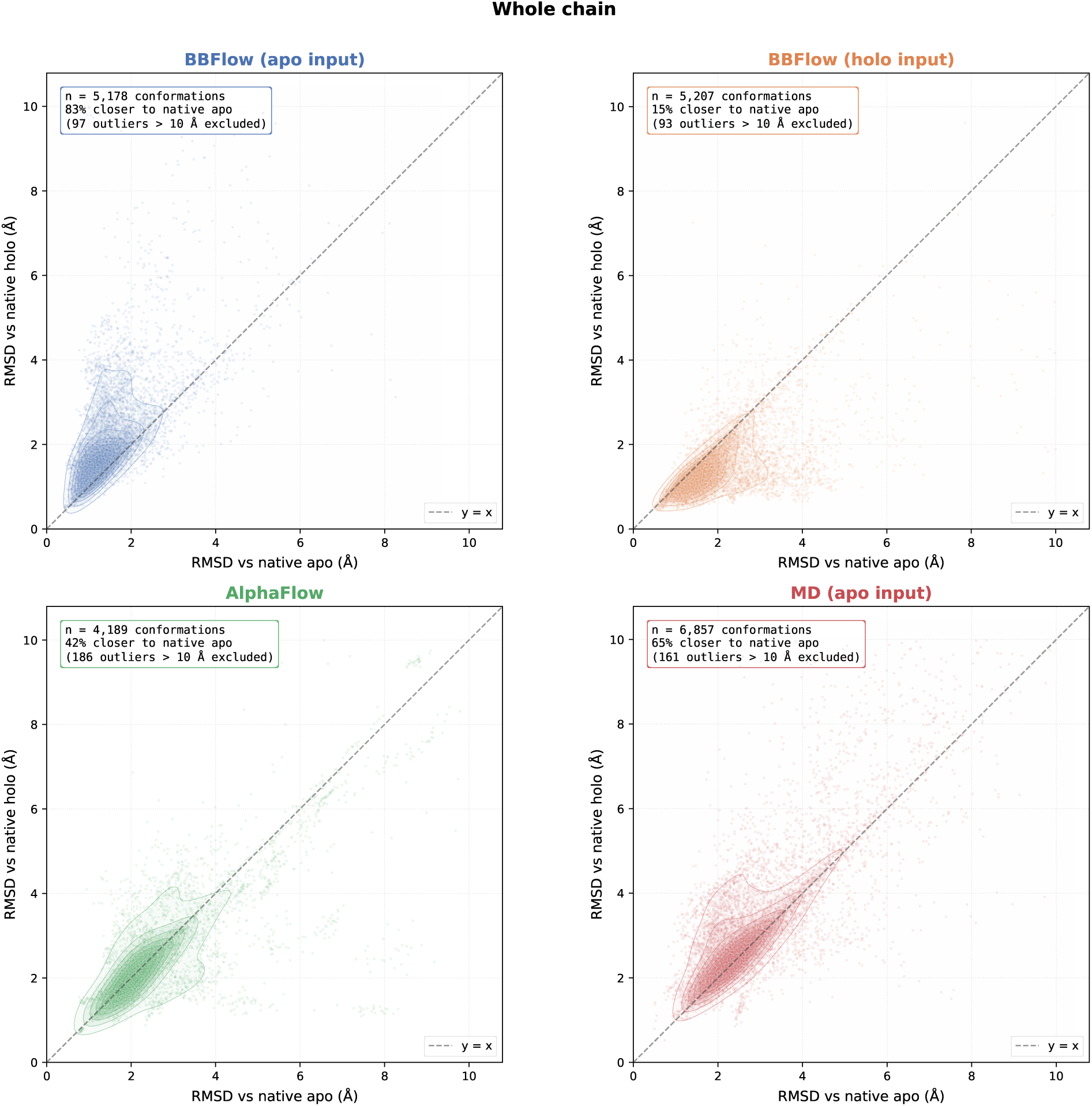
Global conformational similarity of generated ensembles to native states

**Figure S6:**
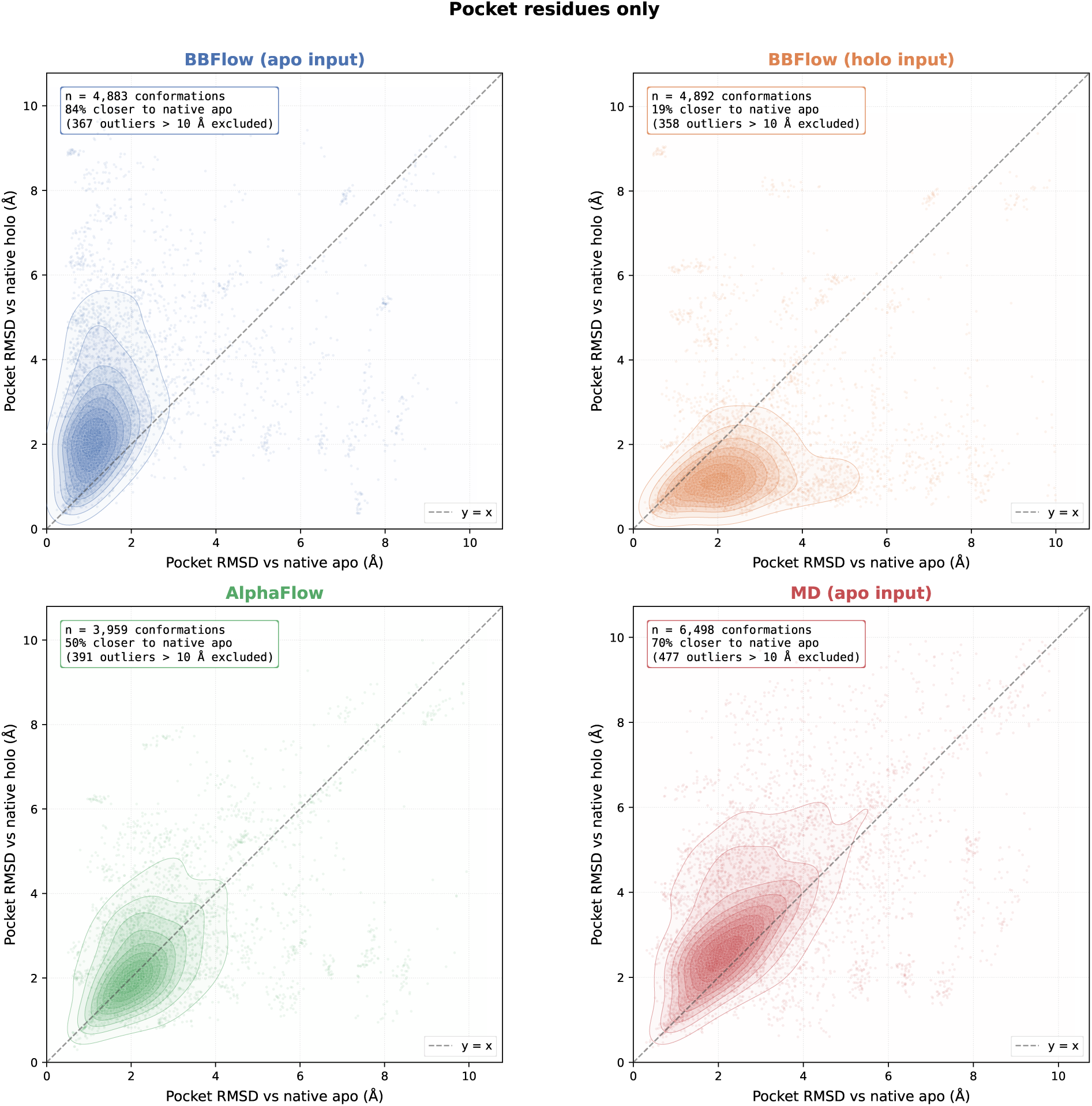
Pocket-only conformational similarity of generated ensembles to native states

**Figure S7:**
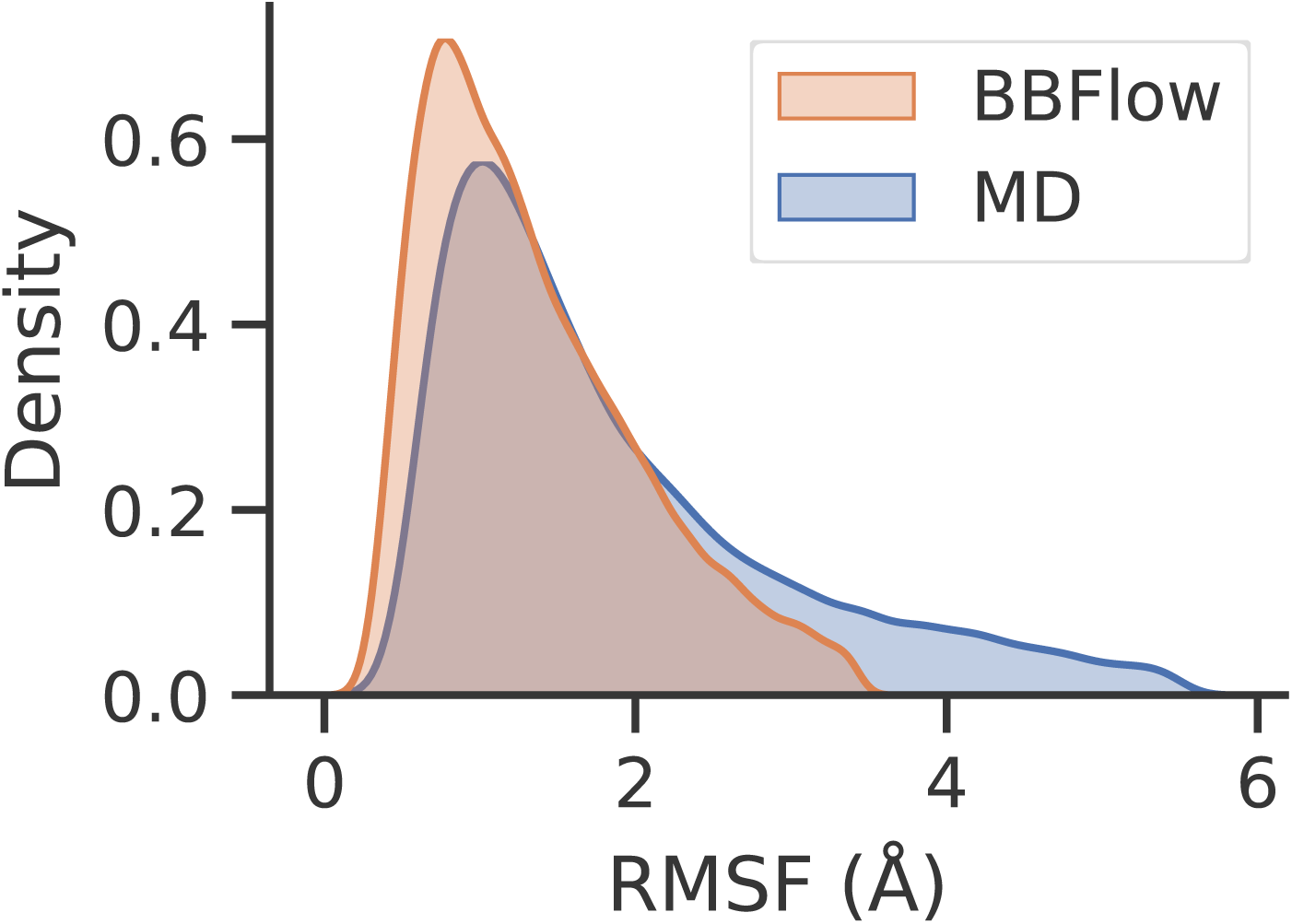
Distribution of per-residue RMSF values from MD simulations and BBflow-generated ensembles across common cryptobench entries. Outliers (*>* 99^th^ percentile) removed.

**Figure S8:**
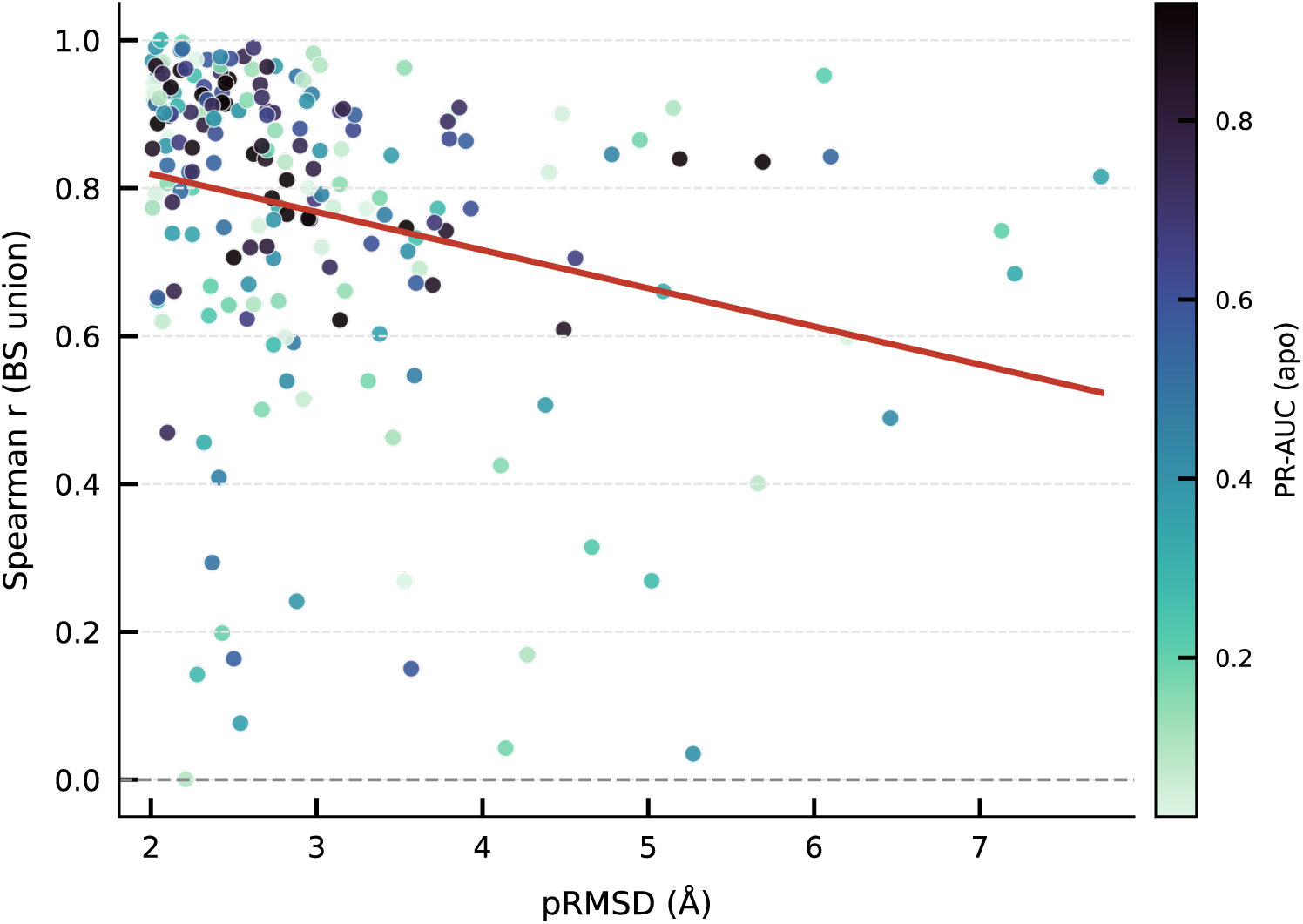
Scatter plot displaying the Spearman correlation (*r*) of residue-level predictions across the union of apo and holo binding site (BS) residues against the pocket root-mean-square deviation (pRMSD, Å) after outlier removal. Each point represents an individual target, colored by the PR-AUC metric on the corresponding apo state.

**Figure S9:**
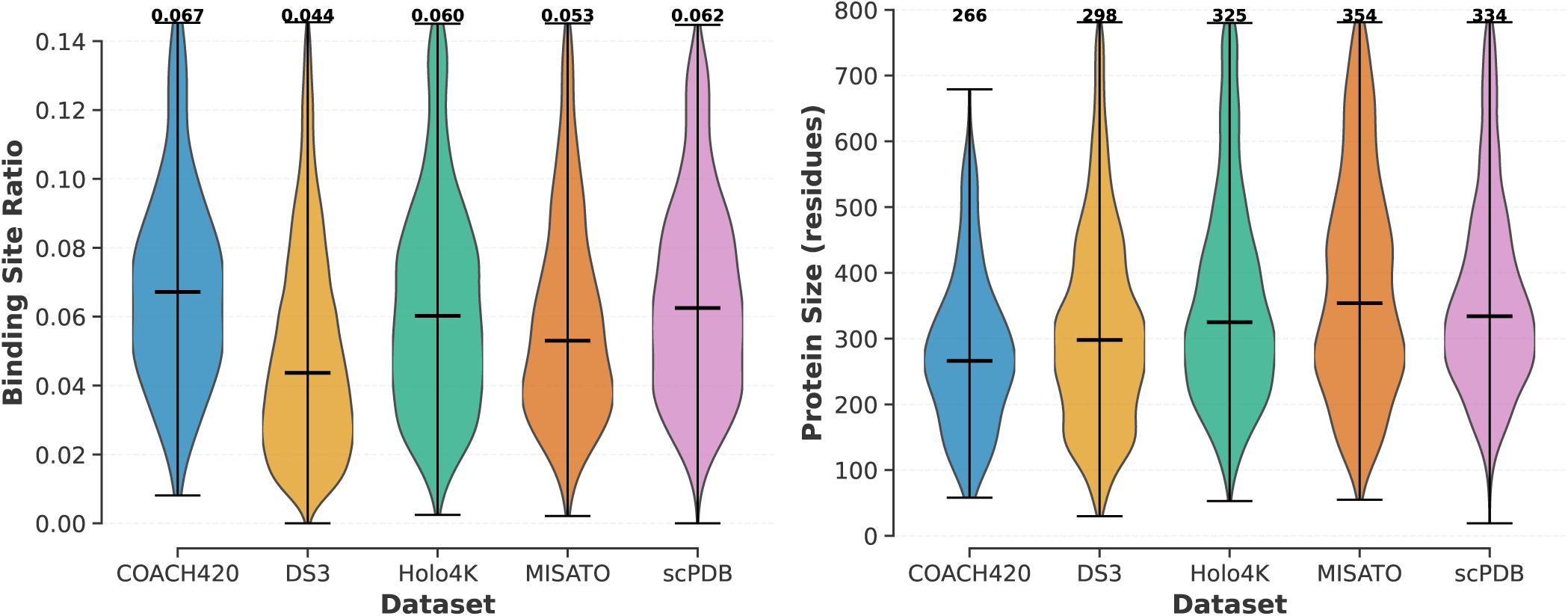
Comparison of binding-site coverage and protein size distributions across datasets after Interquartile Range (IQR)-based outlier filtering.

**Figure S10:**
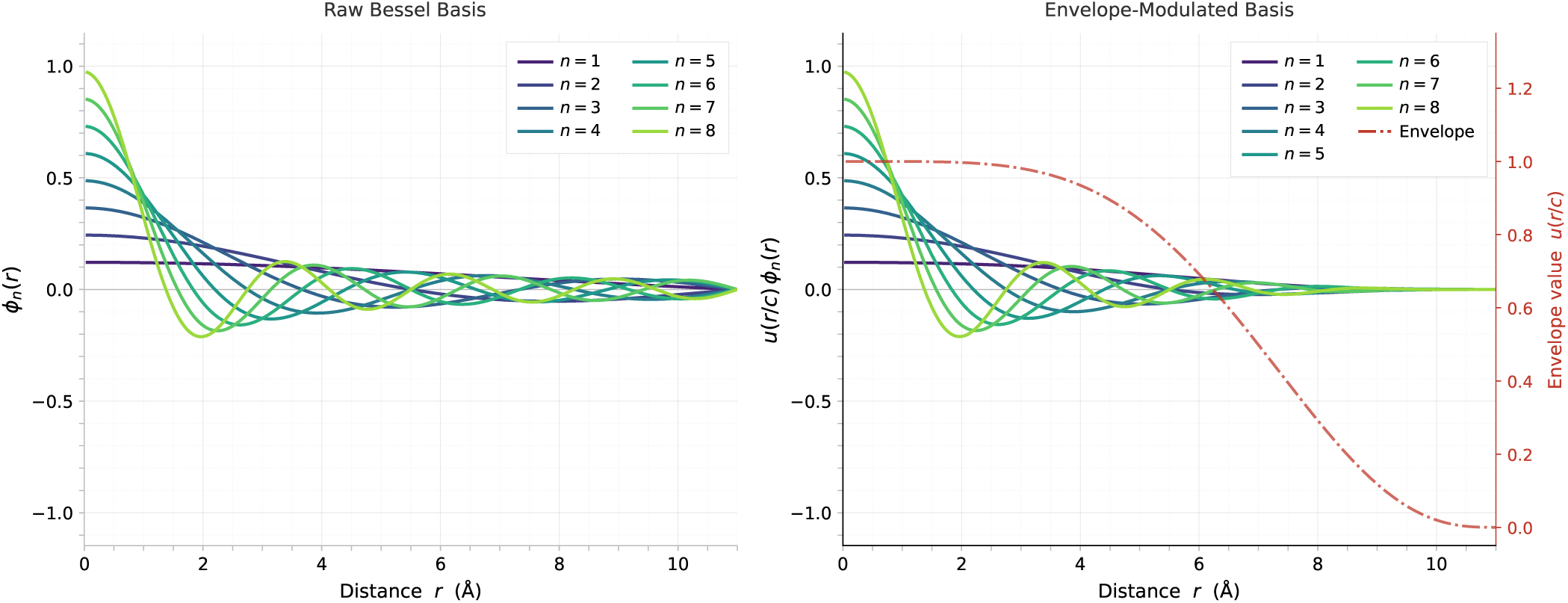
Left: The raw Bessel basis functions *ϕ_n_*(*r*) for *n ∈* {1*, … ,* 8} plotted against distance *r*. Right: The corresponding envelope-modulated basis functions *u*(*r/c*)*ϕ_n_*(*r*), illustrating how the smooth envelope decay function *u*(*r/c*) (dash-dotted red line, right axis) enforces a soft cutoff constraint at larger distances.

**Figure S11:**
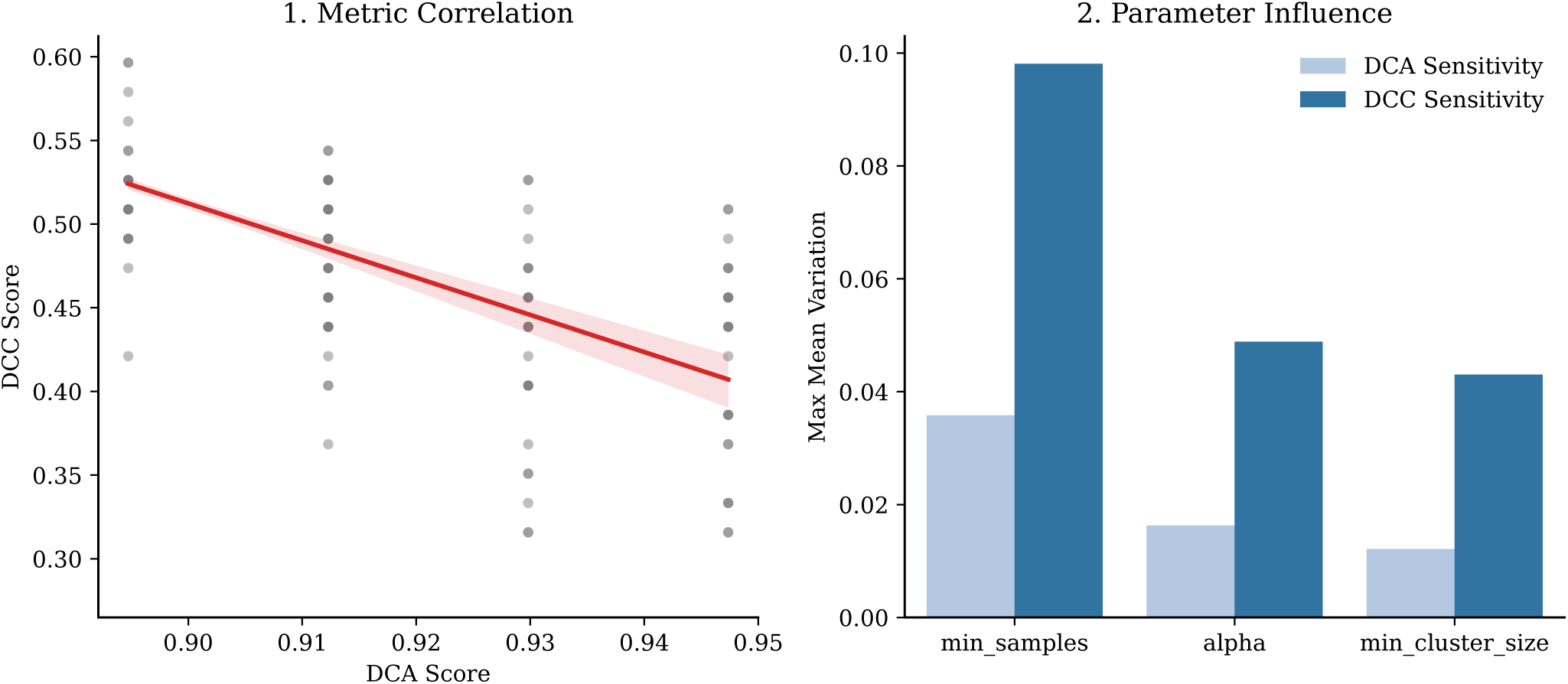
HDBSCAN hyperparameter sensitivity analysis grid search. (1) Inverse linear correlation between DCA and DCC scores. (2) Parameter influence on metric variation. The grid search evaluated min cluster size *∈* {10, 15*, … ,* 40}, min samples *∈* {1, 5*, … ,* 30}, and reweighting coefficient *α ∈* {0.1, 0.5*, … ,* 3.0}. Model selection was performed jointly, identifying min samples as the dominant driver of variance for both metrics.

## 8.2 Supplementary Tables

**Table S1:**
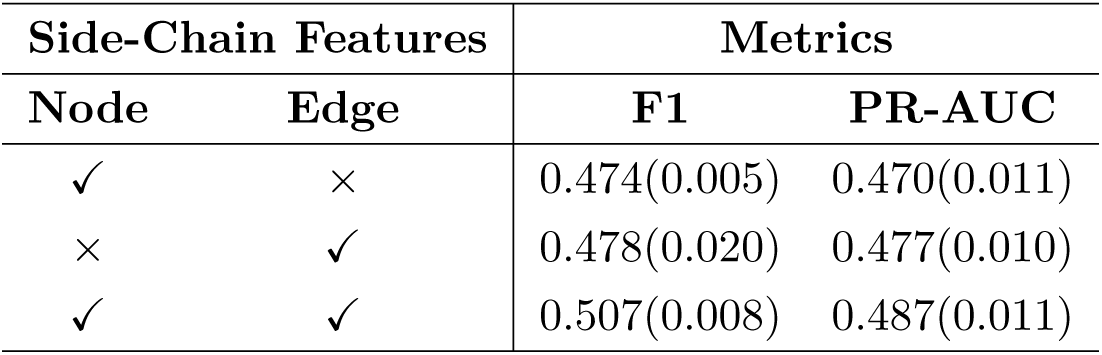
Ablation analysis of explicit side-chain orientation features at node and edge levels.

| Side-Chain Features |  | Metrics |  |
| --- | --- | --- | --- |
| Node | Edge | F1 | PR-AUC |
| ✓ | × | 0.474(0.005) | 0.470(0.011) |
| × | ✓ | 0.478(0.020) | 0.477(0.010) |
| ✓ | ✓ | 0.507(0.008) | 0.487(0.011) |

**Table S2:** Effect of increasing spherical harmonic and embedding irrep orders on validation performance across the MISATO and AlphaFlow datasets.

| SH Irreps | Emb. Irreps | F1 | PR-AUC |
| --- | --- | --- | --- |
| *Dataset: MISATO* |  |  |  |
| $\leq 1$ | $\leq 1$ | 0.4497 | 0.4542 |
| $\leq 1$ | $\leq 2$ | 0.4811 | 0.4877 |
| $\leq 1$ | $\leq 3$ | 0.4969 | 0.5031 |
| $\leq 1$ | $\leq 4$ | 0.4812 | 0.4922 |
| $\leq 2$ | $\leq 2$ | **0.5108** | **0.5095** |
| $\leq 2$ | $\leq 3$ | 0.4997 | 0.5088 |
| $\leq 2$ | $\leq 4$ | 0.5024 | 0.5030 |
| *Dataset: AlphaFlow* |  |  |  |
| $\leq 1$ | $\leq 3$ | 0.5153 | 0.5577 |
| $\leq 2$ | $\leq 2$ | **0.5270** | **0.5752** |
| $\leq 2$ | $\leq 3$ | 0.5193 | 0.5692 |

**Table S3:** Original size of the datasets and number of entries in the successfully generated ensembles. COACH64 contains 64 binding sites corresponding to 59 unique PDB structures.

| **Dataset** | **Original** | **Ensembles** |
| --- | --- | --- |
| scPDB | 5020 | 4997 |
| COACH(mlig) | 300 | 297 |
| HOLO4K(mlig) | 3448 | 3339 |
| DS3(train+valid) | 29689 | 25613 |
| DS3(test) | 1081 | 954 |
| COACH64<sup>1</sup> | 59 | 57 |
| CryptoBench(train) | 885 | 849 |
| CryptoBench(test) | 222 | 212 |

**Table S4:**
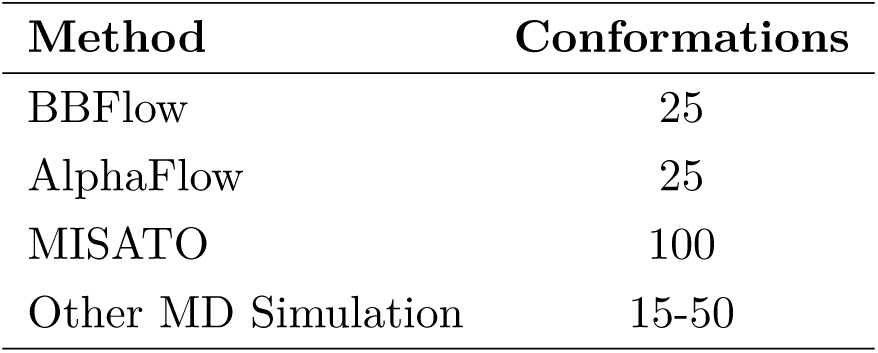
Number of conformations for each protein per method.

| **Method** | **Conformations** |
| --- | --- |
| BBFlow | 25 |
| AlphaFlow | 25 |
| MISATO | 100 |
| Other MD Simulation | 15-50 |

**Table S5:** Training and model hyperparameters, and computational resources.

| Hyperparameter | Value |
| --- | --- |
| Optimizer | AdamW |
| Focal Loss - $\alpha$ | 0.75 |
| Focal Loss - $\gamma$ | 2 |
| Number of epochs | 8 |
| Log Steps | 512 |
| Scheduler update | every log step |
| Warmup steps | 2 |
| Early stopping patience steps | 40 log step |
| Maximum learning rate | $8 \times 10^{-4}$ |
| Minimum learning rate | $5 \times 10^{-6}$ |
| Batch size (trained on complexes) | 2 |
| Batch size (trained on single chains) | 3 |
| Number of Layers | 4 |
| Node-level MOR | [(128, 0), (64, 1), (32, 2)] |
| Attention head MOR | [(16, 0), (8, 1), (4, 2)] |
| Number of attention heads | 4 |
| BesselBasisLayer - Cutoff radius ($\text{\AA}$) | 11 |
| BesselBasisLayer - hidden dim | 128 |
| BesselBasisLayer - envelope exponent | 5 |
| Feed forward network - Dropout rate | 0.1 |
| HDBSCAN - $\alpha$ | 1.5 |
| HDBSCAN - Minimum Cluster Size | 25 |
| HDBSCAN - Minimum Samples | 10 |
| HDBSCAN - Binary Cutoff | 0.4 |
| Hardware | 1 × NVIDIA L40S (45 GiB) |
| Encoder training time | ∼12 hours |
| Set aggregation training time | ∼4 hours |

